# The native glycocalyx is an ordered, self-assembled hierarchical micro- and nanoarray lamellar structure conserved in evolution

**DOI:** 10.1101/2024.07.15.603518

**Authors:** Emanuela Garbarino, Guruprakash Subbiahdoss, Andrea Scheberl, Erik Reimhult, Gradimir Misevic

## Abstract

The native ultrastructure of the glycocalyx remained unknown despite its functional importance in cellular recognition/adhesion and selective filtration. The major components of this universal extracellular coat, mucins, proteoglycans, glyconectins, and hyaluronan, share similar physicochemical properties of high molecular weight, glycan richness, and amply hydrated bottlebrush polymer morphologies with comparable intramolecular anionic charge distribution. The diversity of these glycoconjugate intermolecular binding under physiologically highly hydrated and specific ionic conditions keeps the native glycocalyx structure enabling it to function. Irrespective of the intricacy of the glycocalyx physiological milieu preservation and molecular organization, only a dehydrated non-native state presenting an artefactual unorganized fiber mesh was imaged. Using cryo-SEM after cryo-preservation with minimal sublimation to conserve water, ion distribution, and the native intermolecular interactions, we unveil well-organized lamellae of glycoconjugates self-assembled in hierarchical micro- and nanoarrays for the glycocalyx of human cell and self-assembled glyconectin glycocalyx from an evolutionary most distant sponge despite differences in sequence and composition. Our combined AFM binding strength measurements and cryo-SEM imply that evolutionarily preserved glycocalyx micro- and nano-morphologies are formed by thermodynamically driven self-assembly of glycoconjugates having similar physico-chemical properties.

**Teaser:** The extracellular glycocalyx coat is a self-organizing ultrastructure in human and sponge cells.

## Introduction

The glycocalyx is a glycan-rich, thick coating of all cell surfaces where the plasma membrane encounters the environment (Kuo, Gandhi, Zia, & Paszek, 2018; G. Misevic & Garbarino, 2021). It is often more narrowly referred to as the extracellular mucus layer of the apical portion of epithelial or endothelial cells (Kesimer et al., 2013). Glycocalyx glycoconjugates, such as mucins, proteoglycans, hyaluronan, and glyconectins, play an essential role in the initial step of all interactions with the surrounding, like self-non-self-recognition and adhesion, as well as in the selective binding and filtering of ions, smaller organic compounds, and biopolymers (Bansil, Stanley, & Thomas LaMont, 1995; Buffone & Weaver, 2020; Carpenter et al., 2021; Cummings, 2019; Dammer et al., 1995; Ebong, MacAluso, Spray, & Tarbell, 2011; Kesimer et al., 2013; G. Misevic & Garbarino, 2021; G N Misevic, Finne, & Burger, 1987; Möckl, 2020; Popescu & Misevic, 1997; Sheng & Hasnain, 2022; Simpson, Schaefer, Hascall, & Esko, 2022; Thornton, Rousseau, & McGuckin, 2008; Thornton & Sheehan, 2004; Varki, 2011, 2017; Varki, A., Cummings, R. D., Esko, J. D., Freeze, H. H., Stanley, P., Bertozzi, C. R., Hart, G. W., Etzler, 2022; Zeng, 2017). Their functions closely link to their structures, which are controlled by a balance of entropic hydration forces and local polymeric glycan and protein segment interactions for highly hydrated, multimillion Dalton molecular weight, amply charged, and flexible bottlebrush-like glycoconjugate polymers.

Many studies have focused on inter- and intra-molecular binding of the glycocalyx components via protein-protein, protein-glycan, and glycan-glycan interactions and their relation to the glycocalyx functions. However, there are abundant examples of supramolecular structures determining critical biological functions. It is an open question if the biochemically diverse glycocalyx molecules yield morphologically similar cellular coats. The conserved physicochemical properties, morphological and topological molecular architectures, and binding strength energetics of the diverse types of glycocalyx glycoconjugates from different evolutionary branches imply similar structural roles and even the potential for thermodynamically driven self-assembly of similar ultrastructures. Hence, it is critical to visualize the glycocalyx ultrastructure to improve our understanding of its multitude of functional properties.

Cryo-scanning electron microscopy (cryo-SEM) is the most suitable tool for visualizing the ultrastructure of biological samples in their native form. Ideally, it involves cryo-preservation via high-pressure freezing followed by freeze-fracture and minimal sublimation prior to imaging using SEM under cryo-conditions. Only with this approach can the delicate ultrastructure of the glycocalyx be maintained and visualized.

Unfortunately, the glycocalyx has only been imaged using classical SEM or cryo-SEM after destructive sample preparations involving either chemical fixation and ethanol dehydration or cryo-preservation, including fixation and almost complete sublimation prior to imaging (Critchfield et al., 2013; Jongebloed, Stokroos, Van Der Want, & Kalicharan, 1999; Koga, Kusumi, Bochimoto, Watanabe, & Ushiki, 2015; Li et al., 2021; Sun et al., 2020; Takano, Maekawa, & Takamizawa, 1979; Valk et al., 2002). The expected results of such harsh treatments are drastic chemical and conformational changes in the glycoconjugates. The altered biochemical and biophysical properties of chemically fixed glycocalyx glycoconjugate polymers overcome native molecular interactions. Similarly, removing water and ions during critical point drying or deep etching severely affects the interaction energetics.

These changes will inevitably result in a non-native artefactual ultrastructure appearance. Thus, the glycocalyx has been visualized as a mesh of thick fibers with macromolecularly large pores, lacking higher-order organization (2, 21–30).

## Results

### Preserving the native glycocalyx ultrastructure

To investigate the potential shortcomings of earlier approaches, we examined the level of organization of the glycocalyx ultrastructure closest to the native state by high-pressure freezing 210 MPa, followed by freeze-fracture and a minimal sublimation step (etching), shorter than 120 s at -105 °C and 5.6×10^−7^ mbar, and subsequently coated with ∼1 nm of platinum (see Materials and Methods). This approach strongly reduces artifacts. It maintains the hydrated ion micro-environment around the hydrophilic and highly charged glycoconjugate biopolymers. Hence, we preserve the native polymer conformations and intermolecular bindings essential for sustaining the physiological glycocalyx ultrastructure (Dammer et al., 1995).

We used two mammalian tumor cell lines, MDA-MB-231 (human breast carcinoma) and HT-29 (human colon adenocarcinoma), as models. These mucous glycocalyx-producing cells have been used to study several functions and molecular structures of mucins, proteoglycans, and other glycoconjugates (Gowda, Bhavanandan, & Davidson, 1986; C. Huet, Sahuquillo-Merino, Coudrier, & Louvard, 1987; G. Huet et al., 1995; Maoret et al., 1989; Niv, Byrd, Ho, Dahiya, & Kim, 1992; Soncin, Shapiro, & Fett, 1994). Furthermore, we investigated the ultrastructure of in vitro self-assembled glyconectin 2 (GN 2) from the Porifera *Halichondria panicea*, glycocalyx as the model system of one of the simplest and most evolutionary distant multicellular organisms to mammals. Glyconectins are the major component of the Porifera glycocalyx and are essential in Porifera’s self-non-self-recognition and adhesion via glycan-glycan interactions (Dammer et al., 1995; Guerardel et al., 2004; G. Misevic & Garbarino, 2021; G N Misevic & Burger, 1993; G N Misevic et al., 1987; Gradimir N. Misevic et al., 2004; Popescu & Misevic, 1997). The glyconectin family is also an abundant component of the human gut glycocalyx epithelium and Echinodermata embryonal glycocalyx (G. Misevic, Checiu, & Popescu, 2021; G. Misevic & Garbarino, 2021; Gradimir N. Misevic & Popescu, 1995).

### Organized lamellar ultrastructure of glycocalyx in mammalian cells

The glycocalyx ultrastructure of mammalian HT29 cells directly grown on carriers and examined by cryo-SEM after cryo-preservation and minimal sublimation for 20 s showed well-organized micro- and nano-arrays of linear structures covering distances of up to several hundred micrometers (Figure 1A-C). The structured areas surrounding the cells are their glycocalyx, containing mucins and proteoglycans (Gowda et al., 1986; C. Huet et al., 1987; G. Huet et al., 1995; Maoret et al., 1989; Niv et al., 1992; Soncin et al., 1994). The absence of chemical or mechanical treatment of the cells and surrounding glycocalyx removed only a superficial layer of water from the outermost features of glycocalyx glycoconjugates on the fractured surface. The physiological water content, ion distribution, and intermolecular interactions were maintained, which enabled imaging of the preserved native ultrastructure. Control cryo-SEM experiments performed only with PBS buffer (under both minimal and high etching) or cell culture media containing 10% fetal calf serum (FCS) did not show any lamellar structure organization (Figure S1), thereby excluding artifacts due to salt precipitation during sublimation to forming lamellar structures in cryo-SEM (Liang et al., 2022; Liang, Teng, Chou, & Libera, 2017). Hence, the highly organized lamellae represent the native glycocalyx structure.

**Fig. 1.**
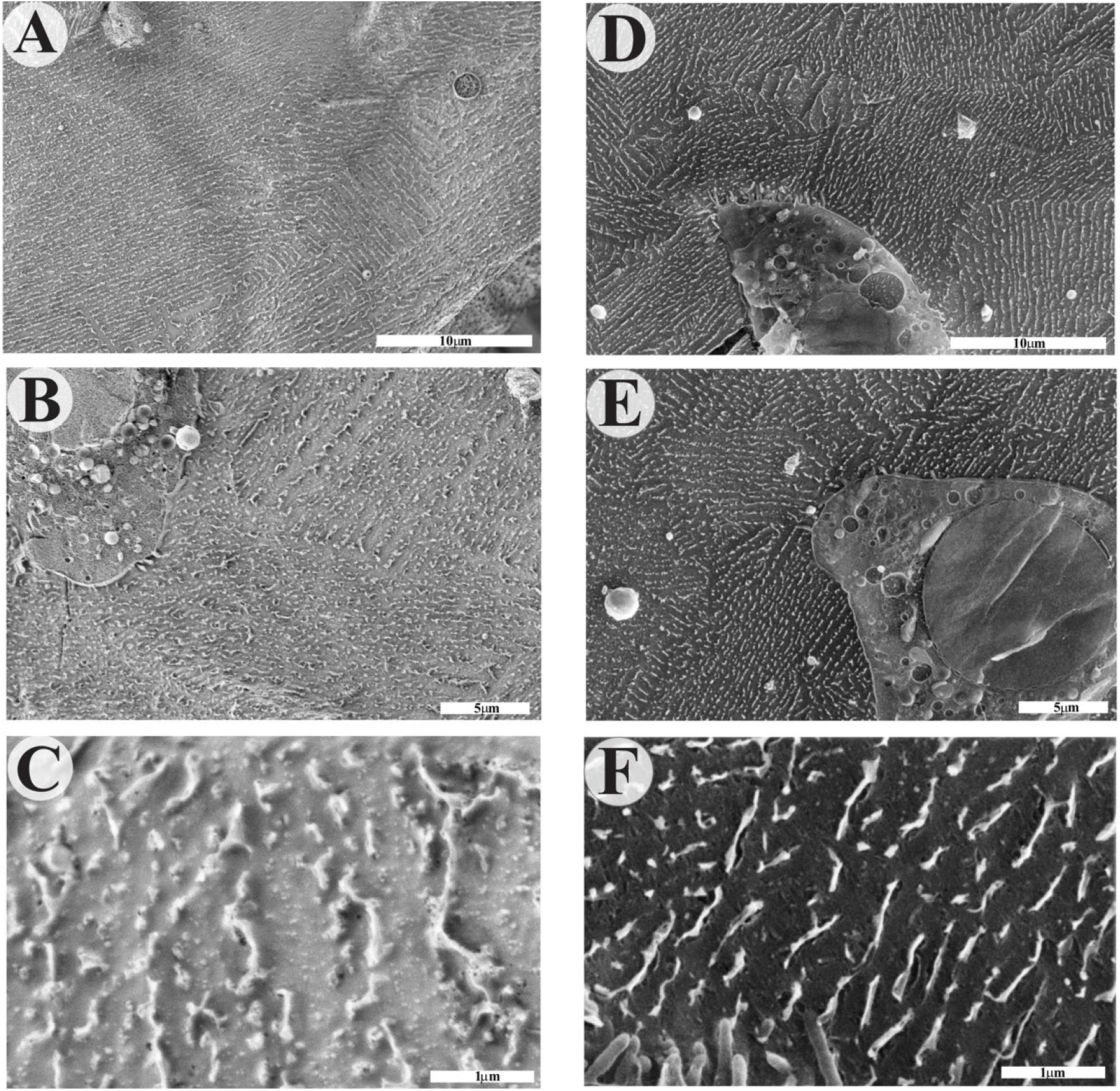
Cryo-SEM of HT-29 cells glycocalyx. (**A**-**C**) Imaging was performed on HT-29 cells grown directly on 3 mm carriers. (**D**-**F**) Imaging was performed on HT-29 cells grown on cell culture Petri dishes after gentle scraping, followed by transfer to carriers. All samples (**A**-**F**) were high-pressure frozen, freeze-fractured at -120 °C, etched at -105 °C, and coated with 4 nm Pt and 4 nm carbon at -120 °C.

The same cryo-preparation and imaging procedure yielded identical lamellar structures for HT-29 cells grown on regular cell culture Petri dishes after gentle scraping and transferring to carriers (Figure 1D-F). Hence, the structures are observed independent of cell preparation, with the scraping approach providing the benefit of reproducibly showing cells together with the glycocalyx in the fracture plane, while fracture planes in the carrier predominantly went through the mucoid glycocalyx.

Therefore, cryo-SEM images of MDA-MB-231 (Figure 2A and B) and HT-29 (Figure 1D-F and 2D and E) cells and their surrounding glycocalyx grown on regular cell culture Petri dishes after gentle scraping and transferring to carriers were compared after 20 s of sublimation. Both cell types showed large extracellular areas of over 1000 μm^2^ covered by a multitude of hierarchically well-organized micro- and nanoarrays of parallel lamellar ultrastructure extending from the cell surface and microvilli, connecting distant cells. The native structure suggests glycan interactions connect the observed proteoglycan lamellae (Figure 2B and E). The highly organized structure contrasts with previous reports of a disorganized, porous mesh of the glycocalyx in HT-29 and other cell types in tissues (Barmpatsalou et al., 2021; Critchfield et al., 2013; C. Huet et al., 1987; Meziu et al., 2021; Sun et al., 2020; Takano et al., 1979; Valk et al., 2002; Vidavsky et al., 2014). Such structures could be reproduced by sublimating water for 6 min (Figure 2C and F). The over-etching resulted in a complete loss of organized ultrastructure. Therefore, we trace the lack of previous recognition of the ultrastructure of the glycocalyx to preparation methods altering their native appearance.

**Fig. 2.**
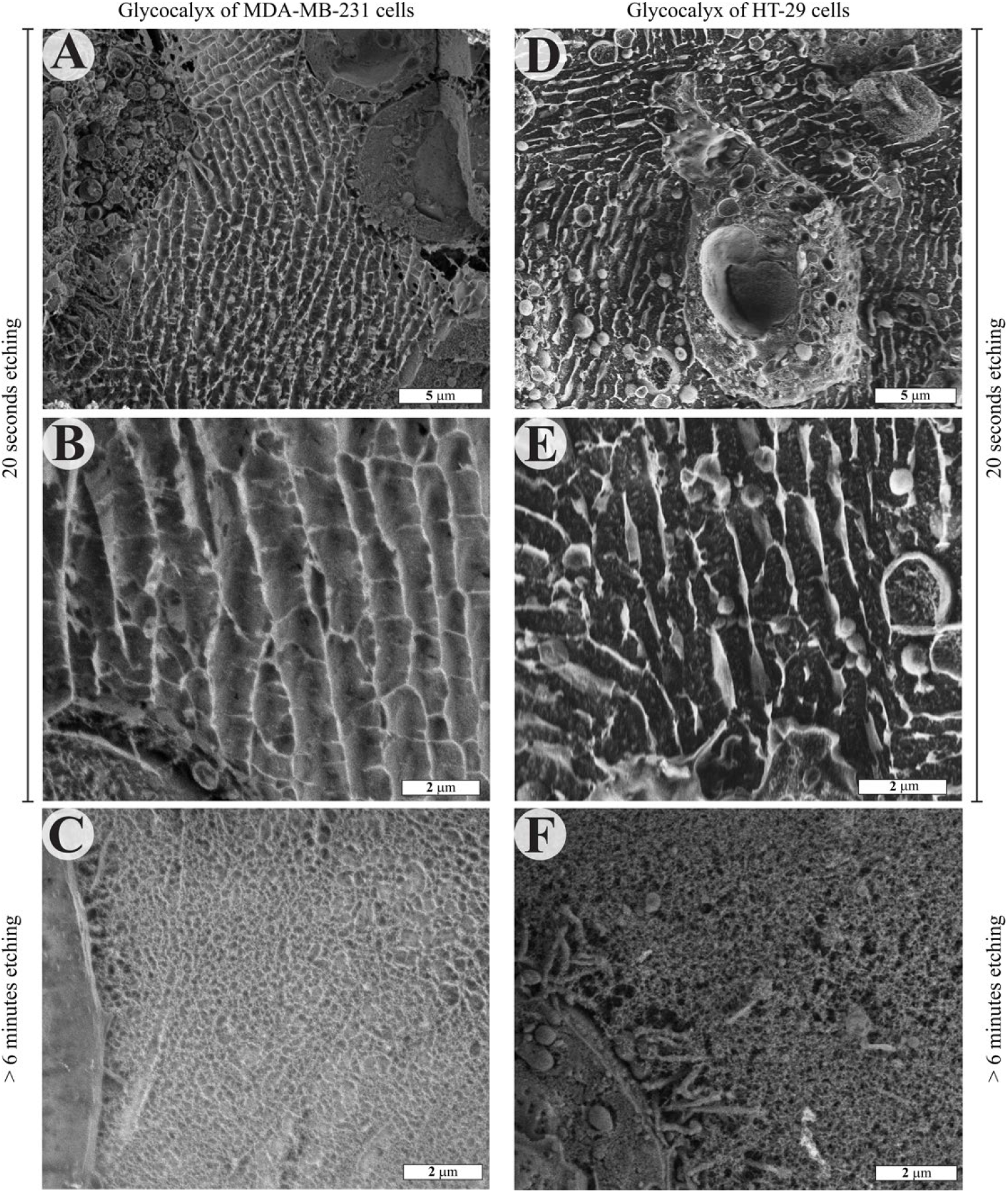
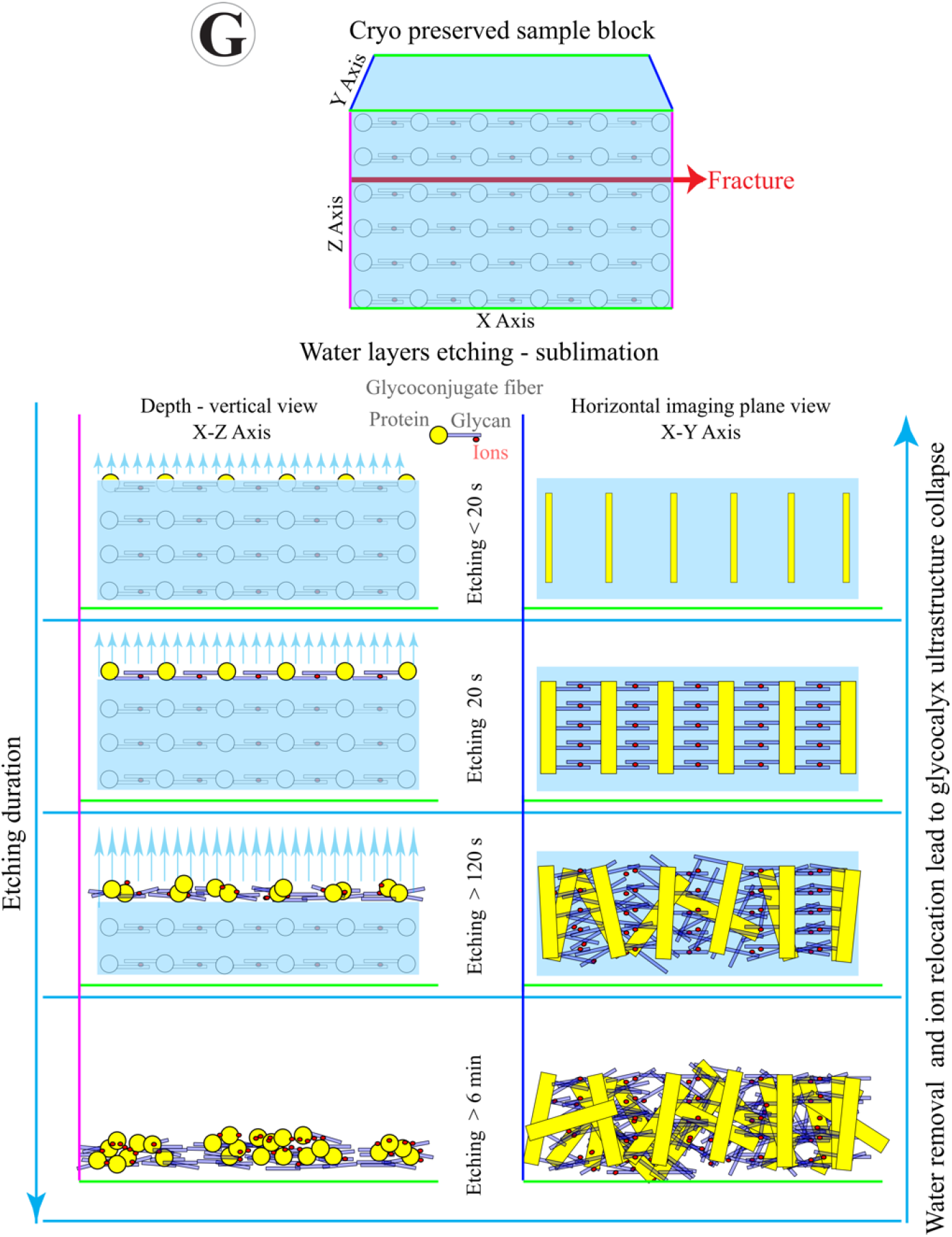
Cryo-SEM of MDA-MB-231 and HT-29 cells glycocalyx with schematics of cryopreservation. The glycocalyx ultrastructure after high-pressure freezing, freeze-fracturing at -110 °C, etching at -105 °C, and ∼1 nm Pt coating at -110 °C. (**A, B**) MDA-MB-231 cells after 20 s of etching. (**C**) MDA-MB-231 cells after 6 min of etching. (**D, E**) HT-29 cells after 20 s of etching. (**F**) HT-29 cells after 6 min of etching. (**G**) Schematic model of the glycocalyx ultrastructure collapse due to water removal and ion relocation.

Deep etching completely removes interstitial water, which is at over 70% of the sample’s mass, thus leaving empty intermolecular space and forcing a relocation of ions to proteins and glycans (Figure 2G). Hydration contributes the dominant part of glycopolymer energetics, determining the conformation. Therefore, water removal results in a drastic conformational change and breaking of the ionic inter-glycan bonds, leading to the collapse of the ultrafine glycan structures into thicker and less organized fibers (Figure 2C and F).

### Organized lamellar ultrastructure of Porifera glyconectin self-assembled glycocalyx

The highly organized micro- and nano-array of the glycocalyx structure does not originate directly from the cells, as shown in Figure 2. This suggests that self-assembling secreted glycoconjugates formed the ordered network. Therefore, we investigated if glyconectins could spontaneously arrange in such well-organized ultrastructures by self-assembly in vitro. We used purified Porifera glyconectin 2 as a simple glycocalyx model system for which we could easily obtain glyconectins in large amounts (see Material and Methods) (Guerardel et al., 2004; Gradimir N. Misevic et al., 2004). GN 2, self-assembled directly on carriers at physiological 10 mM CaCl2 concentration (Guerardel et al., 2004; Gradimir N. Misevic et al., 2004), was prepared for cryo-SEM imaging using the cryo-preservation, fracturing, and etching procedure described for the human MDA-MB-231 and HT-29 cell lines. After 20 s or 100 s of etching at -105 °C and 5.6×10^−7^ mbar, cryo-SEM images showed that GN 2 assembled into a well-organized ultrastructure of lamellae comprised of GN 2 nanofibers covering the entire sample area (>1000 μm^2^) (Figure 3 A and B). The observed ultrastructure resembles the mammalian glycocalyx of both MDA-MB-231 and HT-29 cells in Figure 2A, B, D, and E. The experiment with prolonged etching over several minutes of cryo-fractured GN 2 resulted in a collapse of the ultrastructure, yielding a disorganized, porous mesh (Figure 3C), as for the mammalian cell lines (Figure 2C and F). Hence, this class of glyconectins can reproduce the dominant features of the glycocalyx organization of tissues purely by self-assembly without active cellular processes.

**Fig. 3.**
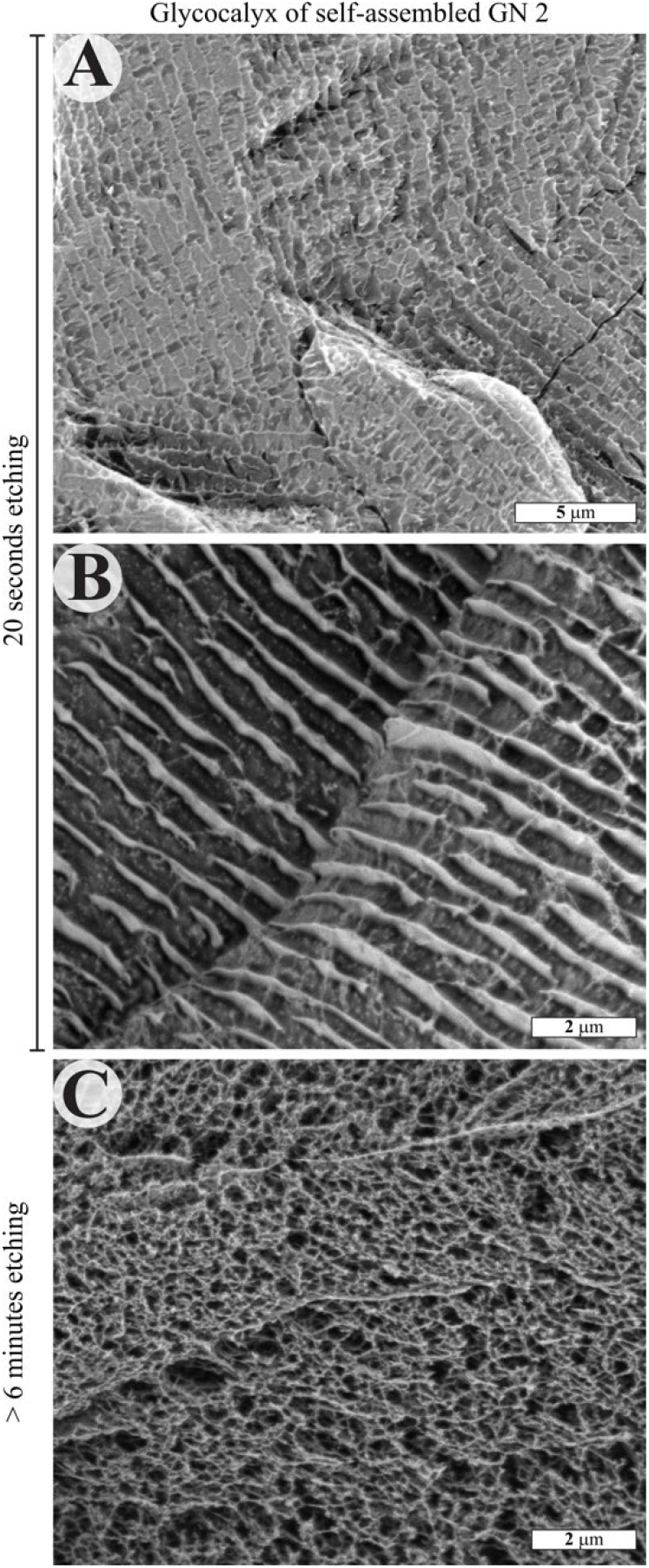
Cryo-SEM of self-assembled GN 2 glycocalyx. The ultrastructure of self-assembled GN 2 obtained from 1mg/ml GN 2 in 0.5 M NaCl, 10 mM HEPES pH 7.4, 2 mM CaCl2, and 0.01 % NaN3, adjusted to physiological 10 mM CaCl2 was imaged by cryo-SEM after high-pressure freezing, freeze-fracturing at -110 °C, etching at -105 °C, and ∼1 nm Pt coating at -110 °C. (**A**) after 20 s of etching. (**B**) after 100 s of etching. (**C**) after 6 min of etching.

Self-assembly was only observed in the presence of calcium ions. To further demonstrate that the structures depended on Ca^2+^-mediated glycan-glycan bonds, we removed calcium ions from a self-assembled GN 2 glycocalyx by briefly incubating it in 50 mM EDTA, which resulted in the dissociation of the lamellar ultrastructure (Fig. S2). The dissociated architecture revealed single, roughly equidistantly spaced and clustered GN 2, which had lost their lamellar order but were still in their native, fully hydrated, and now mutually repelling conformation and structure.

### Model of Porifera glyconectin glycocalyx ultrastructure

Atomic force microscopy (AFM) imaging in the air of GN 2 adsorbed on mica revealed 500 to 660 nm long glyconectin 1-like features, comprising globular subunits of the protein core, each ∼80 nm in diameter, with 10 to 12 glycan side chains arranged in a bottle-brush morphology appearing as 300 to 360 nm long structures (Figure 4D). Similar morphologies can be observed in Figure 4A and B, which provides high-resolution cryo-SEM images of the structure of GN 2 after 60 s and in Figure 3C after 100 s etching. Thus, it exposes superficial parts of the glyconectins. These images suggest a hierarchical micro- and nanoarray of lamellae form from repeating units of self-assembled GN 2 with the lamellae separated by 300-650 nm.

**Fig. 4.**
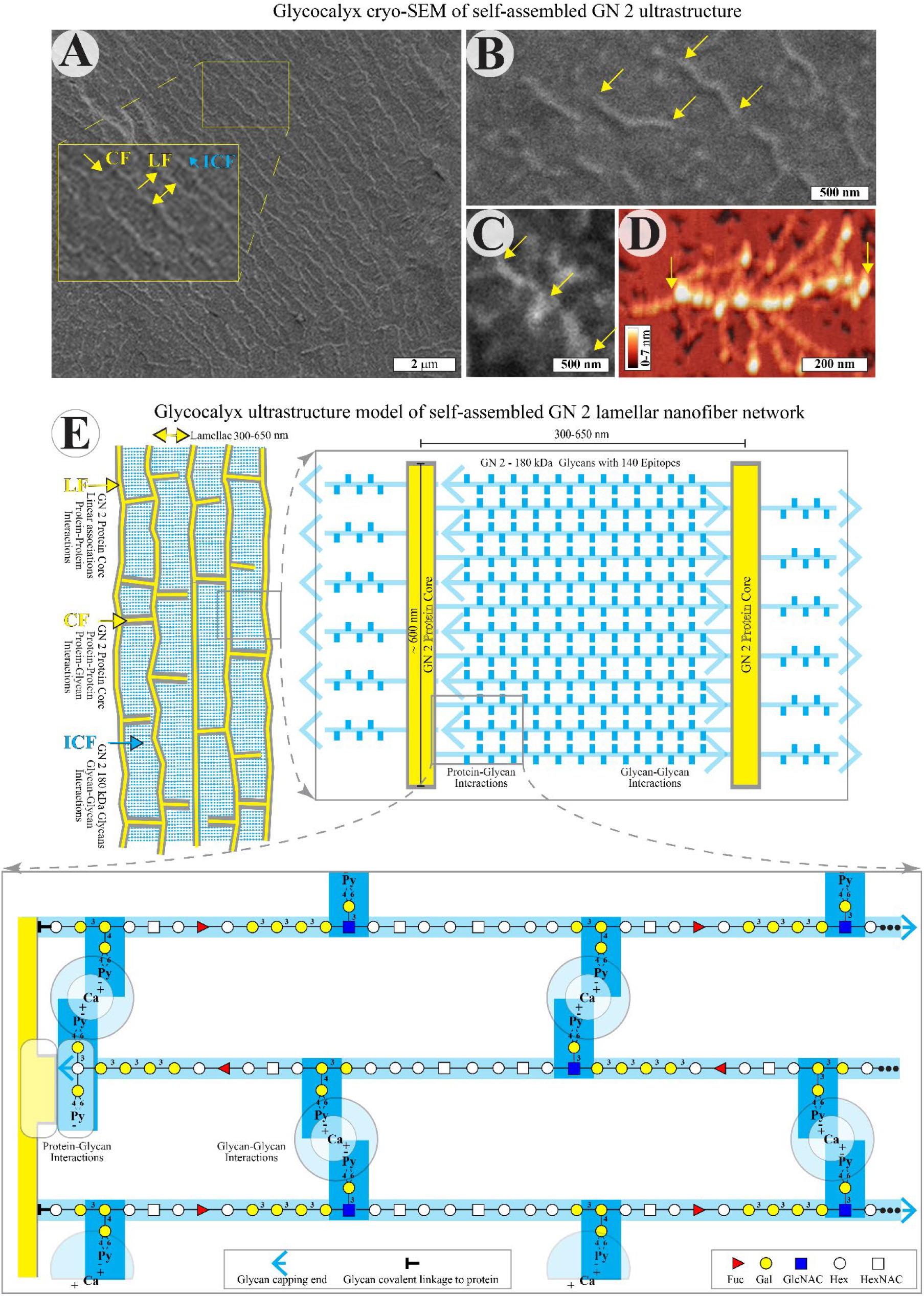
Cryo-SEM, AFM, and a model of self-assembled GN 2 glycocalyx. The ultrastructure of a GN 2 gel obtained from 1mg/ml GN 2 in 0.5 M NaCl, 10 mM HEPES pH 7.4, 2 mM CaCl2, and 0.01 % NaN3, adjusted to physiological 10 mM CaCl2 was imaged by cryo-SEM after high-pressure freezing, freeze-fracturing at - 110 °C, etching at -105 °C, and ∼1 nm Pt coating at -110 °C. (**A, B**) after 60 s of etching. (**C**) after 100 s of etching. (**D**) AFM of a GN 2 molecule spread and dried on mica. (**E**) Schematic model of GN 2 self-assembled glycocalyx ultrastructure based on AFM, cryo-SEM, and sequencing of 180 kDa GN 2 glycans. LF indicates long fibers, CF indicates connecting fibers, and ICF indicates inter-connecting fibers. Fibers are presented in geometrically rigid form. The number of epitopes represented as blue vertical lines of GN 2 180 kDa glycans are enlarged, and their number is 10 times reduced for visualization purposes. The structural model of reconstituted GN 2 glycocalyx is based on partial sequencing of GN 2 glycans and biochemical analyses.

We propose the structural model in Figure 4E based on the molecular morphology of GN 2 obtained from our cryo-SEM and AFM results and previous reports on the structure and function of isolated GN (Guerardel et al., 2004; G. Misevic & Garbarino, 2021; Gradimir N. Misevic et al., 2004). We identify the Long Fibers (LFs) at the edges of lamellae as protein cores linked by protein-protein and/or protein-glycan binding (Figure 4A). Connecting Fibers (CFs) are individual GN 2, linking LFs by protein-protein and/or protein-glycan binding (Figure 3A). We suggest the Inter-Connected Fibers (ICFs) of GN 2 are 180 kDa glycans connecting GN 2 molecules through homophilic glycan-glycan Ca^2+^-dependent binding.

This agrees with previous reports revealing that the purified GN 2 is the major glycocalyx component in Porifera *Halichondria panicea* with Mr = 10 ×10^6^ Da and 21% of its mass comprising of 10 to 12 copies of the large acidic glycans of Mr = 180 kDa (Guerardel et al., 2004; Gradimir N. Misevic et al., 2004). GN 2 was partially sequenced and functionally characterized as a cell recognition and adhesion molecule functioning via Ca^2+^-dependent self-association that is mediated by a highly polyvalent glycan-glycan binding of pyruvylated Py(4,6)Gal(1–4)Gal and Py(4,6)Gal(1–3)GlcNAc epitopes, each estimated with ∼70 copies per glycan molecule and ∼800 copies per GN 2 (Figure 4E) (Guerardel et al., 2004; Gradimir N. Misevic et al., 2004). Such, in sum, strong but locally reversible polyvalent binding between the 180 kDa glycans self-organizes the GN 2 LF and CF protein cores, forming a highly regular skeleton network. The combination of protein nanofiber lamellae and interstitial reversible glycan interactions enable both recognition and filtration functions with the possibility of reorganizing the matrix by local interactions or environmental changes.

Based on the comparable sizes and physico-chemical properties of glyconectins, proteoglycans, mucins, and hyaluronan, we propose a similar glycocalyx model to explain our observations of the two cell lines’ glycocalyxes.

### Measurements of forces between a single pair of glyconectin 2

To investigate the molecular mechanism and the intermolecular forces driving the self-assembly of GN 2 and keeping the ultrastructure integrity, we used AFM to measure the rupture forces between individual pairs of GN 2 under native (seawater) physiological conditions. GN 2 was covalently attached to gold-coated mica and cantilever tips via a self-assembled monolayer using a previously published method (Dammer et al., 1995; Gradimir N. Misevic, Karamanos, & Misevic, 2009) (Figure 5A). We chose the cantilever tip size to accommodate only one GN 2 molecule; this and the low density of GN 2 crosslinked to the flat surfaces assured interactions between single molecular pairs. The average homophilic binding force between a single pair of GN 2 in seawater containing the physiological 10 mM Ca^2+^ for the marine Porifera was 150 pN, and the maximum was about 300 pN (Figure 5B). The Ca^2+^-induced binding was demonstrated by no attraction between the GN 2 (rare events of < 10 pN) at 1 mM Ca^2+^ (Figure 5C), i.e., the retraction curve retraces the approach curves of both Ca^2+^ concentrations. Selectivity for Ca^2+^ was confirmed by the inability of 10 mM Mg^2+^ to restore GN 2 binding (Figure 5D).

**Fig. 5.**
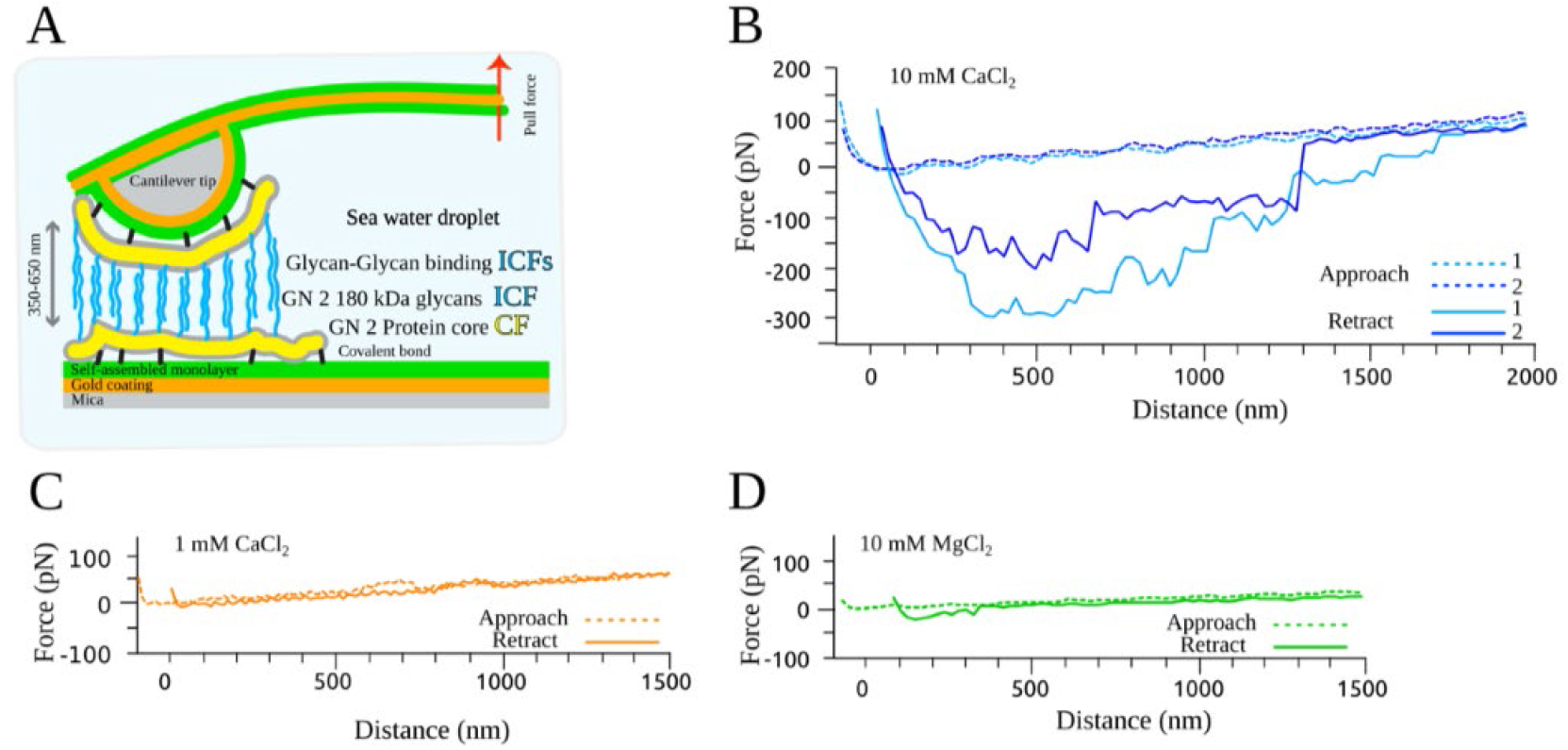
AFM measurements of force between a single pair of GN 2 molecules. (**A**) Schematic representation of AFM intermolecular force measurements between GN 2 covalently attached to a planar surface and an AFM cantilever tip. (**B**) Force-distance measurements between two GN 2 molecules in a droplet of seawater with 10 mM Ca^2+^. Two typical approach-retract cycles, light blue lines represent cycle 1 and dark blue lines represent cycle 2, dashed lines indicate approach, and solid lines indicate retract parts of cycles. (**C**) A force approach-retract cycle between two GN 2 molecules in a droplet of seawater with 1 mM Ca^2+^. (**D**) A force approach-retract cycle between two GN 2 molecules in a droplet of seawater with 10 mM Mg^2+^.

The shape of an approach-retract cycle (binding-unbinding) in physiological seawater (10 mM Ca^2+^) revealed multiple unbinding events, occurring either gradually or stepwise at separation distances ranging from 350 nm up to 1750 nm (Figure 5B). The maximal binding forces of about 300 pN per a single GN 2 pair were reached at distances between 350 nm and 650 nm. The maximum unbinding force distances correlate well with the LF-LF (GN 2 protein core) distances observed in Figure 3. GN 2 has 10 to 12 copies of 180 kDa glycan chains. We get an upper bound on the involved forces by assuming all chains were involved in the GN 2 pair binding, i.e., ∼30 pN per single glycan chain.

The binding involves polyvalent weak interactions. It allows highly dynamic binding, frequent rearrangement, and reforming of bonds at different intermolecular distances, glycan chain stretching, angles, and arrangements. The measured unbinding events, therefore, reflect different lengths of chain binding rather than the unbinding of all different GN 2 chains. It explains the broad distribution of unbinding forces and the occasional observation of gradual (slip) unbinding. Assuming a lamellar arrangement indicated by Figures 1-4, each GN 2 will form bonds to at least four other GN 2 and fully saturate its potential for glycan-glycan binding and achieve a binding force of more than 300 pN in the ordered self-assembled structure. The combination of single-chain, homophilic interactions in the low pN range and overall GN 2 interactions close to the nN range are suitable for self-assembling stable structures that can be highly dynamic on the nanoscale. Dynamic rearrangement is a prerequisite for self-assembly, as observed in Figures 2 and 3. The combination of dynamic local interaction and high overall macromolecular affinities can lead to ordered, stable ultrastructures with characteristic size, topologies, and intermolecular distances. Re-arrangement necessary for biological functions can be facilitated globally by altering the ionic composition or pH of the environment or locally by substituting the glycan-glycan interactions of individual chains by inserting replacement interactions.

## Discussion

Cryo-SEM is the ultimate approach for studying the delicate and highly hydrated glycocalyx ultrastructure at the supramolecular level. However, it has not been properly applied to imaging of the glycocalyx, resulting in a loss of ordered structures (Barmpatsalou et al., 2021; Meziu et al., 2021; Sun et al., 2020; Takano et al., 1979; Valk et al., 2002; Vidavsky et al., 2014). We designed a protocol without fixation but with minimal sublimation to conserve water and ion distribution between glycans to investigate the hypothesis that glycocalyx molecules can self-assemble driven by ion-mediated self-recognition and polymer hydration. By doing so, we could demonstrate a highly organized ultrastructure in the glycocalyx and reproduce previous observations of a disorganized mesh by continuing the sublimation of stabilizing amorphous ice (water). AFM force-distance measurements of GN 2 binding interactions showed the plausibility of supramolecular arrangement resulting from the magnitude of individual and collective unbinding events. We measured forces between individual Ca^2+^-mediated pyruvylated sugars on the level of hydrogen bonds and inter-glycan chain (ICF) forces comparable to forces between short DNA strands, indicating the ability to dynamically self-assemble these segments. However, the aggregate forces holding one GN 2 in the superstructure are on the order of multiple biotin-avidin bonds, providing long-term stability to the overall structure. A well-defined force minimum suggests a preferred GN 2 protein core distance correlating with the ultrastructure observed by non-destructive cryo-SEM.

The observed similarities of the native glycocalyx ultrastructure for different mammalian cell types and the self-assembled glycocalyx of the evolutionary very distant Porifera with different mucins, proteoglycans, hyaluronan, and/or glyconectins may be unexpected at first glance. However, these macromolecules share comparable physico-chemical properties, e.g., very high molecular weight, bottle-brush polymer morphology, high hydration, net-negative charge, and ion-mediated bonding patterns. Several hundred negative charges carried by sialic acid in mucins, uronic acids in hyaluronan, and sulfated monosaccharides, uronic acid, and pyruvylated monosaccharides in proteoglycans and glyconectins are similarly distributed in these molecules. These dense charge patterns with similar density and intramolecular distribution within the topology of hydrated bottle-brush polymeric structures infer highly polyvalent electrostatic intermolecular interactions under physiological conditions of salt concentration in water. Functional homologs with structural but no sequence similarities from evolutionary well-separated organisms are known. They can perform similar functions but with different specific interactions and be tailored to various environments. S-layer proteins are one such example, controlling the interaction of large classes of archaea and prokaryotes with their environment (Sleytr, Schuster, Egelseer, & Pum, 2014).

Besides previously described protein-glycan and protein-protein binding related to the glycocalyx, our prior data (Dammer et al., 1995; Guerardel et al., 2004; G. Misevic & Garbarino, 2021; G N Misevic & Burger, 1993; G N Misevic et al., 1987; Gradimir N. Misevic et al., 2004; Popescu & Misevic, 1997) and here presented findings also unveil polyvalent ionic and Ca^+2^ dependent glycan-glycan interactions in GN 2 as the essential driving force for self-assembling of glycocalyx into an organized ultrastructure. Direct evidence of glycan-glycan associations in the glycocalyx was shown for glyconectin glycans in Porifera (Dammer et al., 1995; Guerardel et al., 2004; G. Misevic & Garbarino, 2021; G N Misevic & Burger, 1993; G N Misevic et al., 1987; Gradimir N. Misevic et al., 2004; Popescu & Misevic, 1997) and Echinodermata (G. Misevic et al., 2021), and for hyaluronan gelation (Garg & Hales, 2004).

The ordered self-assembled morphology of the glycocalyx at the micro and nano levels enables tuning of biological material properties and functions, such as viscoelasticity, water and ion retention, and specificity and selectivity of filtration at the hierarchical levels of ions, macromolecules, viruses, bacteria, and cells. Hence, our findings open new avenues of scientific interrogation, implying the importance of structure-function relationships on the supramolecular level and deciphering the functions of extracellular matrices. It could greatly influence the understanding of selective multimolecular self-assembly in biopolymer research and self-recognition at epithelial and endothelial glycocalyx layers in biology and medicine.

## Materials and Methods

### Cell culture and harvesting

Human breast carcinoma (MDA-MB-231) and human colon adenocarcinoma (HT-29) cells, obtained from German collection of Microorganisms and Cell Culture GmbH, Braunschweig, Germany.

Prior to the seeding of HT29 cells, carriers were first sterilized with 70 % ethanol and then placed into cell culture Petri dishes. Subsequently, carriers were washed three times in culture dishes with PBS followed by three washes and 10 min incubation with Dulbecco’s Modified Eagle’s Medium (DMEM) with low glucose, supplemented with 10% fetal bovine serum (FBS) and 20 mM HEPES at 37 ºC in a humidified 5% CO2 atmosphere, After aspiration of the medium, HT-29 cells were seeded in cell culture Petri dishes containing carriers and were grown in Dulbecco’s Modified Eagle’s Medium (DMEM) with low glucose, supplemented with 10% FBS and 20 mM HEPES at 37 ºC in a humidified 5% CO2 atmosphere. After reaching 85-90% of confluence, dishes were washed three times with phosphate buffer saline at pH 7.2 to remove the residual medium. Finally, carriers were taken out of the Petri dishes and closed with the upper carrier part, followed by high-pressure freezing without any mechanical or chemical manipulation of cells, thus completely preserving the native configuration.

Mammalian cells (MDA-MD-231 or HT-29) and glycocalyx were scraped and deposited onto carriers: The cells in T75 culture flasks, after reaching 85-90% of confluency, cells were washed three times with phosphate buffer saline at pH 7.2 (PBS) before harvesting to remove the residual medium. Finally, the whole cellular layer with surrounding glycocalyx was removed from the plates by gentle scraping with a cell scraper. The high viability of scraped cells was directly confirmed with Methylene blue and optical microscopy. The harvested cells were immediately transferred to aluminum- or gold-coated (Type A) specimen carriers (Leica Microsystems Inc., Austria) for high-pressure freezing.

### Glyconectin 2 isolation

*Halichondria panicea* was purchased from the Marine Biological Laboratory in Woods Hole, MA, USA. Sponges were rinsed with Ca^2+^- and Mg^2+^-free seawater (462 mM NaCl, 10.7 mM KCl, 7 mM Na2SO4, and 2.1 mM NaHCO3) (CMF-SW). GN 2 was extracted from fresh cuts of 1-2-centimeter sponge pieces with the CMF-SW at +4 °C for 12 h (Guerardel et al., 2004; Gradimir N. Misevic et al., 2004). Dissociated cells were centrifuged, and the supernatant containing GN 2 was recentrifuged to remove cellular debris as previously described (Guerardel et al., 2004; Gradimir N. Misevic et al., 2004). The supernatant containing GN 2 was adjusted to 20 mM CaCl2 by adding an appropriate volume of 1 M CaCl2. Complete GN 2 gelation occurred after 12 hours of incubation at +4 °C. Gels were centrifuged and subsequently dissolved in 0.5 M NaCl, 10 mM HEPES pH 7.4, containing 2 mM CaCl2 and 0.01 % NaN3. GN 2 was further purified by ultracentrifugation and finally stored in 0.5 M NaCl, 10 mM HEPES pH 7.4, containing 2 mM CaCl2 and 0.01 % NaN3 (Guerardel et al., 2004; Gradimir N. Misevic et al., 2004).

### High-pressure freezing, freeze-fracture, minimal sublimation, and cryo-SEM

High-pressure freezing, freeze-fracture, and cryo-scanning electron microscopy (cryo-SEM) were performed to examine the ultrastructure of MDA-MB-231 and HT-29 glycocalyx in their native forms and of purified glycocalyx component GN 2 of the marine sponge *Halichondria panicea* in its reconstituted gel form. GN 2 was obtained from 1mg/ml of 0.5 M NaCl, 10 mM HEPES pH 7.4, 2 mM CaCl2, and 0.01 % NaN3 adjusted to physiological 10 mM CaCl2. Samples of GN2 or scraped cells were placed between two 3 mm diameter aluminum- or gold-coated (Type A) specimen carriers with 100 µm indentations (Leica Microsystems Inc., Austria). HT29 cells directly grown on the same type of 3 mm diameter aluminum- or gold-coated (Type A) specimen carriers were closed with the same size upper carrier. Assembled carriers were placed between two cylinders into sample cartridges and frozen immediately using a high-pressure freezing machine (Leica EM HPM100, Leica Microsystems Inc., Austria).

The internally integrated flow channels in the cylinders directed the liquid nitrogen (LN2) onto the samples. The air-driven intensifier in HPM100 pressurized the LN2 to 210 MPa and injected it into the freezing chamber. The frozen sample carriers were then mounted onto a sample holder under LN2 and transferred to the Leica EM ACE900 freeze-fracture system (Leica Microsystems Inc., Austria) via a load lock system. This was followed by fracturing at -110 °C using a flat edge knife and sublimation at -105 °C and 5.6×10^−7^ mbar (Leica EM ACE900) for 20 s, 60 s, 100 s, or 6 min for different samples as specified in the text. Next, the samples were e-beam coated with a platinum layer of either 1.1 nm thickness or 4 nm platinum and 4 nm carbon thickness at -110 °C for better contrast and to reduce beam damage to the samples during imaging. Subsequently, the samples were transferred to the SEM via a cryo-transfer system (Leica EM VCT500). Imaging was performed using an Apreo VS SEM (Thermo Scientific, The Netherlands) at an acceleration voltage of 500 V or 1 kV. High-vacuum back-scattered and secondary electrons were detected using two in-lens detectors, T1 and T2, respectively. During SEM imaging, the chamber temperature was maintained at -100 °C and pressure at 10^−4^ Pa, a temperature at which the surface was stable, showing neither ice condensation nor sublimation of water vapor, while at lower temperatures, ice condensation was observed.

### AFM measurements of force between a single pair of GN 2

Freshly cleaved mica and cantilever tips from Park Scientific Instruments, Mountain Views, CA, (0.01 N/m with 200 nm diameter) were coated with 10 nm gold as described previously (Dammer et al., 1995; Gradimir N. Misevic et al., 2009) were incubated with freshly prepared 11-thioundodecanol in analytical-grade methanol for 12 h at room temperature to form a self-assembled monolayer. After washing with methanol 2 times, 50 mg/mL of freshly prepared carbonyldiimidazole in analytical grade methanol was added and incubated for 20 min at room temperature. Mica and cantilever tips were washed with methanol and seawater buffered with 20 mM HEPES, and crosslinking of GN 2 was performed at a concentration of 20 μg/ml CMF-SW with 2 mM Ca^2+^ buffered with 20 mM HEPES at pH 7.4 for 1 h at room temperature. After 5 washes with the same buffer and mounting of functionalized mica and cantilever, AFM measurements (NanoScope Ill, Digital Instruments, Santa Barbara, CA) were performed in a droplet of CMF-SW buffered with 20 mM HEPES at pH 7.4 in the presence of 10 mM Ca^2+^, 2 mM Ca^2+^, and 10 mM Mg^2+^ at room temperature with 0.01–1 Hz on 5–10 different locations. About twenty to forty approach-and-retract cycles over two hours were recorded.

## Supporting information

Supplemental Materials Garbarino et al.

## Acknowledgments

We thank Octavian Popescu for helping with AFM measurements and Andreas Weber for helping with the cell cultures.

## Author contributions

Conceptualization: GM, EG

Methodology: SG, AS, EG, GM, ER

Investigation: EG, SG, GM, AS, ER

Visualization: SG, EG, GM, AS, ER

Supervision: GM, ER, SG

Writing—original draft: EG, GM

Writing—review & editing: EG, GM, SG, ER

## Data and materials availability

All data are available in the main text or the supplementary materials.

## Supplementary Materials

Included in a separate document.

