## Supplemental Materials Garbarino et al. for "The native glycocalyx is an ordered, self-assembled hierarchical micro- and nanoarray lamellar structure conserved in evolution"

Emanuela Garbarino *et al.*

*.*

*Gradimir Misevic.

**This PDF file includes:**

Supplementary Text

Figs. S1 to S2

Supplementary Text

**Results**

**Cryo-SEM of growth medium and PBS**.

Cryo-SEM imaging of the control samples comprising either cell culture growth medium

containing 10% FCS or PBS buffer, cryo-preserved in the same way as cells and GN 2, did not

show any lamellar structure (Figure S1). Grow medium containing 10% FCS revealed globular

protein features (Figure S1A). PBS samples that were more extensively etched to examine

possible salt-related artifacts were flat without structural features (Figure S1B). These results

confirmed that the cryo-SEM visualized lamellar micro and nano-arrays organized ultrastructure

in cryo-preserved cells or GN 2 represent self-assembled glycocalyx glycoconjugates and not

artifacts from residual serum contaminants and/or salts.

**Disassembling ultrastructure of GN 2 self-assembled glycocalyx by removal of Ca^2+^.**

Removing calcium ions from self-assembled GM 2 glycocalyx by EDTA treatment resulted in the dissociation of the lamellar ultrastructure (Figure S2). The non-organized architectural

appearance revealed single GN 2 molecules in their dissociated native state because the natural

environment was still fully hydrated with the physiological concentration and distribution of all

other ions. The highly etched samples presented in Figure 3C were different because they

showed the denatured and collapsed structure due to complete dehydration and relocation of

ions. Furthermore, this data, together with the control experiments presented in Figure S1

showed that well organized lamellae represent self-assembled GN 2 ultrastructures.

**Methods**

**Cryo-SEM of growth medium with 10 % FCS and cryo-SEM of PBS.**

The control samples comprising of either Dulbecco´s Modified Eagle´s Medium (DMEM) with low glucose, supplemented with 10% fetal bovine serum (FBS) and 20 mM HEPES or PBS buffer were placed in an aluminum- or gold-coated (Type A) specimen carrier (Leica Microsystems Inc., Austria) with 100 µm indentations followed by cryo-preservation and processing in the same way as cells and GN 2 using high-pressure freezing, freeze-fracturing at -120 °C, etching at either -110 °C for 20 s or at -95 °C for 10 min, followed by 4 nm Pt and 4 nm carbon coating at -120 °C. Samples carriers were assembled between two cylinders into sample cartridges and were high-pressure frozen using a (Leica EM HPM100 (Leica Microsystems Inc., Austria).

**EDTA treatment of GN 2**.

Self-assembled GN 2 ultrastructure was treated with EDTA directly in an aluminum- or gold-coated (Type A) specimen carrier (Leica Microsystems Inc., Austria) with 100 µm indentations. To 100 µl of 1 mg/ml of GN 2 in 0.5 M NaCl, 2 mM CaCl_2_, 20 mM HEPES pH 7.4, 0.05 % NaN_3_ 5 µl of 0.5 M EDTA in 0.5 M NaCl, 2 mM CaCl_2_, 20 mM HEPES pH 7.4 was added. The sample was incubated in a humid chamber at room temperature for 10 minutes. The second 5 µl aliquot of 0.5 M EDTA in 0.5 M NaCl, 2 mM CaCl_2_, 20 mM HEPES pH 7.4 was added, reaching the final concentration of 50 mM of EDTA in the sample. After 10 minutes of incubation at room temperature, the carrier was closed with the upper carrier part with 100 µm indentations. Then, the sample carriers were assembled between two cylinders into sample cartridges and were high-pressure frozen using a (Leica EM HPM100 (Leica Microsystems Inc., Austria). We used the same procedure of freezing, fracturing, and 20 seconds etching as for not-treated GN 2.

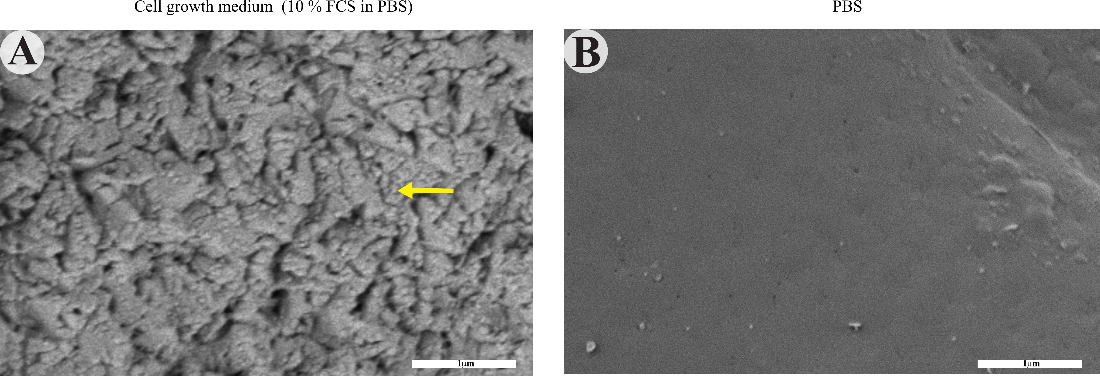

Fig. S1.

**Cryo-SEM of growth medium and PBS control samples.** The control samples cell culture growth medium containing 10% FCS and PBS buffer after high-pressure freezing, freeze-fracturing at -120 °C, etching at -110 °C, and 4 nm Pt and 4 nm carbon coating at -120 °C. (**A**) Cryo-SEM image of cell culture medium containing 10 % FCS after high-pressure freezing, freeze-fracturing at -120 °C, etching at -110 °C for 20 s, and 4 nm Pt and 4 nm carbon coating at -120 °C. The yellow arrow points to a single FCS protein. (**B**) Cryo-SEM of PBS after high-pressure freezing, freeze-fracturing at -120 °C, etching at -95 °C for 10 min, and 4 nm Pt and 4 nm carbon coating at -120 °C.

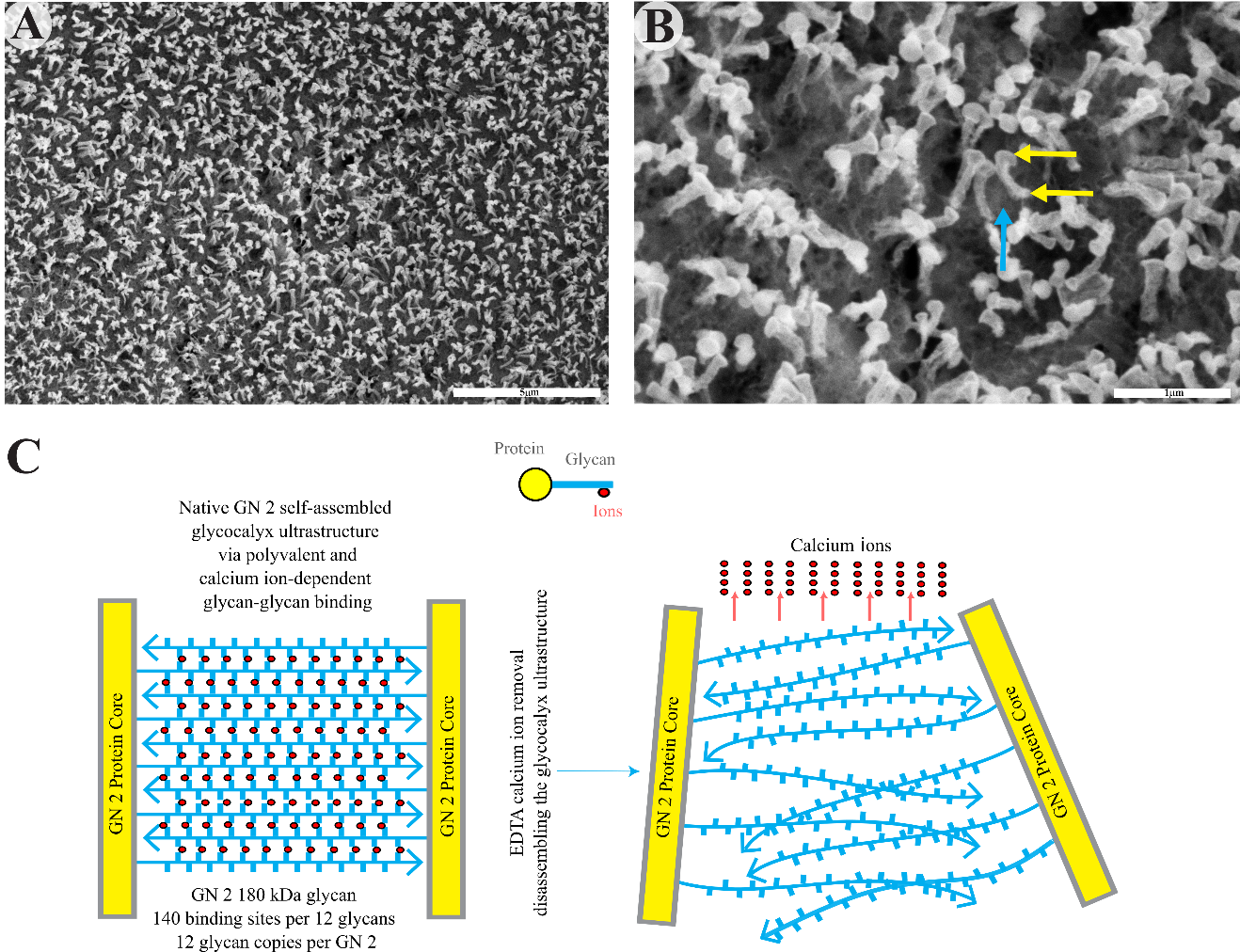

Fig. S2.

**Cryo-SEM of GN 2 ultrastructure disassembled by EDTA treatment.** GN 2 treated with EDTA removing calcium ions mediating the polysaccharide bindings. (**A**) and (**B**) Cryo-SEM after high-pressure freezing, freeze-fracturing at -110 °C, 20 s etching at -105 °C, and 4 nm Pt and 4 nm carbon coating at -110 °C. Two yellow arrows are pointing to the ends of the protein core of a single GN 2 molecule, and the blue arrow points to 180 kDa glycan fibers. (**C**) Schematic model of changes in the GN 2 glycocalyx ultrastructure caused by Ca^2+^ removal.
